# Kinase activity simultaneously determines the constitutive and the orthosteric gating in α_4_β_1/3_δ GABA_A_ receptors in hippocampal granule cells

**DOI:** 10.1101/318543

**Authors:** Nils Ole Dalby, Ulrike Leurs, Christina Birkedahl Falk-Petersen, Petra Scholze, Jacob Krall, Bente Frølund, Petrine Wellendorph

## Abstract

A subset of the GABA_A_ receptors expressed in recombinant systems and neurons is known to exhibit both constitutive- and agonist-induced gating. Two such receptors are the δ-subunit containing GABA_A_ receptors α_4_β_1_δ and α_4_β_3_δ, which are expressed in adult rodent hippocampal dentate gyrus granule cells (DGGCs). Here we show that the GABA_A_ receptor mediated tonic current recorded in the presence of tetrodotoxin in adult rodent DGGCs is almost exclusively mediated by constitutively active δ-subunit containing GABA_A_ receptors and that the constitutive current is absent in recordings at 24 °C or in recordings at 34 °C including an intracellular inhibitor of protein kinase C. These factors simultaneously govern the efficacy of an orthosteric agonist at α_4_β_1/3_δ receptors, Thio-THIP, in a reciprocal manner. In the absence of constitutive receptor activity, the efficacy of Thio-THIP was increased approximately four-fold relative to recording conditions that favors constitutive activity. Further, only under conditions of an absent constitutive current, the classified neutral antagonist gabazine (GBZ) alone, induced a tonic current in DGGCs (EC_50_ 2.1 μM). This effect of GBZ was not seen in recording conditions of high constitutive activity, was inhibited by picrotoxin (PTX), potentiated by DS2, completely absent in δ^-/-^ mice and reduced in β_1_^-/-^ mice, but could not be replicated in human α_4_β_1/3_δ receptors expressed recombinantly in HEK cells. We hypothesize that specific intracellular components in neurons interact with receptors to determine constitutive gating and receptor responsiveness to orthosteric ligands.

**Significance statement:** The presented data highlight how recording conditions for whole cell patch clamp analysis of α_4_β_1/3_δ GABA_A_ receptors can mask important pharmacological effects. Specifically, orthosteric agonists appear with reduced efficacy, and other ligands, here exemplified with the well-known antagonist GBZ, are misinterpreted as being inactive/neutral, although they could have effect in constitutively silent receptors. Unmasking of potential hidden effects are easily done using recording conditions of reduced kinase activity in a relevant neuronal context. It follows that in pathologies with changes in phosphorylation level of δ-subunit containing GABA_A_ receptors, the efficacy of an agonist of these receptors, measured by whole-cell recordings *in vitro*, will not match the efficacy of the same agonist in an unperturbed neuron *in vivo*.

## Introduction

GABA_A_ receptors belong to the Cys-loop superfamily of ion channels. The functional channel that comprises the orthosteric binding site for GABA is a pentamer composed of two α, two β and one γ or δ subunit. Sampled from a population of six α (1-6), three β (1-3), two γ (1-2) and one δ, the possible number of permutations is in the hundreds. However, the existence of assembly rules favoring distinct combinations allow for only a subset to exist *in vivo* (Olsen and Sieghart, 2008; Martenson et al., 2017). The anatomical localization is, with a few exceptions, subunit-specific and typically the γ-subunit containing receptors mediates a synaptic, fast inhibitory post-synaptic current (IPSC), and the δ- and the α_4-6_-subunit containing receptors mediate a slow extrasynaptic tonic current (Farrant and Nusser, 2005; Chandra et al., 2006). The magnitude of the tonic current is region and neuron-specific, and the general view is that synaptic overspill, including release from neurogliaform interneurons, is the main determinant of GABA in the cerebrospinal fluid (CSF) and of the tonic current (reviewed by (Glykys and Mody, 2007; Lee and Maguire, 2014; Overstreet-Wadiche and McBain, 2015). However, several GABA_A_ receptor subtypes in recombinant systems, cultured hippocampal neurons and acute slices also display a degree of constitutive activity (Birnir et al., 2000; McCartney et al., 2007; Tang et al., 2010; Othman et al., 2012; Wlodarczyk et al., 2013; Hoestgaard-Jensen et al., 2014). For example, the receptor subunit combinations α_1/4_β_3_γ_2(L)_ and α_4_β_1/3_δ are capable of both GABA-activated and constitutive gating, the latter (α_4_β_3_δ) in a manner dependent on protein kinase A (PKA) activity (McCartney et al., 2007; Tang et al., 2010; Jensen et al., 2013; Hoestgaard-Jensen et al., 2014). Further, there is evidence to suggest that, while constitutively active, receptor populations of the α_4_β_3_δ subtype in recombinant systems simultaneously display reduced efficacy to low levels of GABA, either by imposing a limitation on the dynamic range for orthosteric agonism or/and by desensitization (Tang et al., 2010; Jensen et al., 2013). The hippocampal dentate gyrus granule cells (DGGCs) in adult rats and mice exhibit near the highest level of co-expressed α_4_, β_1/3_ and δ subunits seen in the brain (Sperk et al., 1997), suggesting that α_4_β_1/3_δ GABA_A_ receptors are very probable inhabitants at peri- and extra-synaptic loci in DGGCs (Wei et al., 2003; Chandra et al., 2006; Zhang et al., 2007; Herd et al., 2008). However, the effect of a selective agonist at α_4_β_1/3_δ receptors suggested, paradoxically, a very limited expression of these receptors in DGGCs, and mainly at synaptic loci (Hoestgaard-Jensen et al., 2014). To examine this problem in detail, the present study was concerned with the role of protein kinase activity on the constitutive current, and further, the relation between constitutive and agonist-mediated gating in DGGCs. Initially we show that the endogenous tonic current recorded in adult rodents DGGCs is predominantly due to constitutive gating and that this channel behavior is determined by recording temperature, intracellular calcium, and by the activity of protein kinase C (PKC). By minimizing the constitutive activity with kinase inhibition, we find that an inverse relation exists between the magnitudes of constitutively active *versus* agonist-induced gating by a subset of δ-subunit containing GABA_A_ receptors in DGGCs. This ability to minimize the constitutive gating by intracellular inhibition of kinase activity has also provided new understanding on the effects of the two different GABA_A_ receptor antagonists bicuculline (BIC) and gabazine (GBZ), because GBZ is assumed to be inactive towards constitutively active receptors (Bai et al., 2001; McCartney et al., 2007; Wlodarczyk et al., 2013). However, under conditions of no apparent constitutive current, we find that GBZ introduces a current in a population of δ-subunit containing GABA_A_ receptors in DGGCs, which is reminiscent of the constitutive current, and accordingly, GBZ is not a neutral antagonist.

## Materials and Methods

### Chemical compounds

4,5,6,7-Tetrahydroisoxazolo[5,4-*c*]pyridin-3-ol (THIP) and 4,5,6,7-tetrahydroisothiazolo[5,4-*c*]pyridin-3-ol (Thio-THIP) were synthesized at the Department of Drug Design and Pharmacology, University of Copenhagen as previously described (Krehan et al., 2003). CGP54626, TTX, SR95531 (gabazine; GBZ), TPMPA, MLA, PKC autoinhibitory peptide (19-36), (4-chloro-[2-(2-thienyl)imadazo[1,2*a*]pyridine-3-yl]benzamide (DS2) and GABA were obtained from Tocris Bioscience (Bristol, UK) and kynurenate, strychnine, atropine, picrotoxin (PTX) and bicuculline-methiodide (BIC) were obtained from Sigma-Aldrich (St. Louis, MO, USA).

### Animal work

Adult male rats were obtained from Envigo and housed in cages holding up to six rats with ad *libitum* access to food and water in rooms maintained at 22-24 °C, 55% humidity at a 12 h/12 h normal night/day cycle.

Similar housing conditions were used for mice. All animal experiments were carried out in accordance with the European Communities Council Directive (86/609/EEC) for the care and use of laboratory animals and the Danish legislation regulating animal experiments.

For mouse studies, δ and β_1_ GABA_A_ receptor subunit null (germline) mutants were employed. A breeding pair of mice heterozygous for the δ deletion (Mihalek et al., 1999) of the strain C57BL/6J ×129Sv/SvJ was kindly provided by Dr. Delia Belelli, University of Dundee, Scotland and bred in-house. Mice carrying deletions in the β_1_ gene were generated and bred in-house as described below. All mice were bred at the specific pathogen-free animal facility at the University of Copenhagen. For both δ and β_1_ deleted strains, heterozygotes were intercrossed to create littermates of all three genotypes (^-/-^, ^+/+^, and ^+/-^) and monitored by PCR genotyping. Only ^-/-^ and ^+/+^ male mice resulting from heterozygous breeding were used for the studies.

### Production of β_1_^-/-^ mice

For the generation of mice with a targeted deletion of *gabrb1*, a Knockout First promoter driven mouse Gabrb1tm1a(KOMP)Wtsi) was purchased from the KOMP repository (www.komp.org) and used to generate a full germline knockout (β_1_^-/-^). To this end, the obtained mouse was first crossed with a global Flp-expressing mouse to generate a conditional *loxP*-flanked (floxed) allele. Secondly, this mouse was crossed with a global Cre-expressing mouse to generate mice with a constitutive knockout allele, β_1_^+/-Cre+^. These were back-crossed into the C57BL/6 background to remove the *Cre* and *Flp* and to produce a genetically pure offspring and carefully monitored by PCR analysis. The resulting mouse lacks exon 4 of the *gabrb1* gene and instead contains one *frt* and one *loxP* site. After several steps of backcrossing to the C57BL/6J background strain, the generation equivalent level was determined by speed congenics using SNP analysis technology (1450 SNP marker panel) (Taconic, Lille Skensved, Denmark). Of seven out of seven tested heterozygous males, all were found to be at least 99.72% identical to the C57BL/6J background, thus at least equivalent to the N8 generation. In general, the mice lacking either one or both deleted alleles (β_1_^+/-^ or β_1_^+/-^) were viable and born at expected Mendelian ratios. The mice were normal in their development, fertile, and did not exhibit obvious behavioral or physical phenotypes. Animals were genotyped using the REDExtract-N-Amp™ Tissue PCR Kit (Sigma Aldrich) according to the manufacturer’s instructions. Primers were designed to discriminate knockout and wildtype β_1_ alleles (forward primer 5’-GTATGGTCAGATGTCCTCAC-3’ and reverse primer 5’-CTGTCTTGCTAGCTTTGAG-3’). The amplification was as follows (94 °C 5 min initial; then 94 °C 30 sec, annealing at 57 °C for 30 sec, extension at 72 °C for 1.5 min for 35 cycles) to generate fragments of 1211 bp for wildtype allele and 572 bp for the knockout allele.

### Slice electrophysiology

Slices of rat brain were obtained from young adult male Sprague Dawley rats weighing 230-300 g at the time of experiment. Mice were 2-4 months old at the time of experiment. At the day of experiment, a single rat or mouse was decapitated and the head immediately immersed in ice-cold artificial CSF (ACSF, composition below) for 6-8 min before dissection of the brain. For dissection and slicing of the brain, ACSF contained (in mM): N-Methyl D-Glucamine (NMDG, 100), NaCl (26), KCl (2.5), CaCl_2_ (1), MgCl_2_ (3), NaHCO_3_ (26), NaH_2_PO_4_ (1.25), D-glucose (10), ascorbate (0.3), pyruvic acid (0.3), kynurenic acid (1), adjusted to 305+/-3 mOsm and aerated with carbogen (95 % O_2_/5 % CO_2_). The brain was glued to the base of a Leica VT1200 S vibratome and cut in 350 μm thick horizontal sections. Slices were stored at 28-29 °C in a carbogenated ACSF of composition as above but without kynurenate and NMDG replaced with equimolar NaCl. Whole cell patch clamp recordings of individual neurons in slices began after a 90 min recovery period, and slices were used up to 4 h thereafter. Recording ACSF were in composition of bulk chemicals similar to storage ACSF, but carried in addition: kynurenate (1 mM), CGP54626 (1 μM), atropine (1 μM), strychnine (1 μM) and TTX (0.5 μM). For electrically evoked IPSCs, TTX was omitted and a Ca^2+^/Mg^2+^ ratio of 2/2 was used. During recordings, slices were maintained in a chamber holding 2 ml of ACSF at a flow rate of 2.8 – 3.0 ml/min at a temperature of 23-24 or 33-34 °C. We used four different conditions for recording of the pharmacological effects of drugs efficacious at neuronal GABA_A_ receptors, which we refer to as condition ***a***, ***b***, ***c*** and ***d***. For condition ***a***, recordings were made at 23-24 °C using an intracellular solution (ICS) composed of (in mM): CsCl (135), NaCl (4), MgCl_2_ (2), N-2-hydroxyethylpiperazine-N’-2-ethanesulphonic acid (HEPES) (10), EGTA (0.05), QX-314 (5), tetraethylammoniumchloride (TEA, 5), Mg-ATP (2), Na_2_-GTP (0.5). The pH of the ICS was adjusted to 7.2 with CsOH and held an osmolarity of 290-295 mOsm. For condition ***b***, recordings were made at 33-34 °C using the same ICS. For condition ***c***, recordings were made at 33-34 °C, using the same ICS but including 10 μM of the auto-inhibitory fragment (19-36) of PKC. For these experiments, an aliquot of the kinase inhibitor peptide was added to a prefiltered (0.22 μm) vial of ICS before use. Condition ***d*** was performed also at 33-34 °C, using ICS of similar composition as in condition ***b***, but in which the concentration of EGTA was increased from 0.05 to 10 mM and CsCl accordingly decreased to maintain osmolarity. Individual neurons were visualized in an IR video-microscope and recorded using thin-walled borosilicate pipettes (Sutter, OD/ID 1.5/1.1 mm) with a 6-8 MΩ resistance (condition ***a***-***d***). To ensure a good diffusion of the ICS into the neuron, we used an 8-9 min period between establishing the whole-cell configuration and beginning recording of baseline activity. Recordings of phasic and tonic currents consisted of a 3 min baseline period, followed by a 6 min drug period (Thio-THIP, THIP and GBZ) and 3-5 min of BIC, GBZ or PTX. Cell capacitance (CP) and series resistance (RS) were measured every 3-4 min throughout the recording and 80 % compensated for RS. Cells were excluded from analysis if values for CP and RS deviated more than 30 % from initial values determined at baseline. For evoked synaptic events, a constant current stimulus isolator (A365, WPI) was used to deliver a 0.5 ms rectangular current pulse via a bipolar theta glass electrode (Sutter, OD 1.5 mm), pulled to obtain 3-5 μm openings and filled with ACSF. Stimulation electrodes were positioned either proximal (maximum 20 μm from cell soma) or distal, in the outermost 20 μm of the molecular layer. Whole-cell recordings were made using a Multiclamp 700A patch clamp amplifier controlled by pClamp 9 software and digitized at 10 kHz with a digidata 1322 (all Molecular Devices) and filtered (8-pole Bessell) at 3 kHz.

### Analysis of patch clamp recordings in DGGCs

The average holding current value was assessed as the center-value, xc, of a single Gaussian fit of the general expression 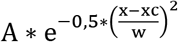 to single-point trace-values in a two-min period (one-min for final antagonist) sampled at a 100 ms interval. The analysis-window for baseline- and drug-period ended immediately prior to beginning of bath perfusion of the ensuing drug. The tonic current density (TCD) was calculated as the difference in holding currents divided by the cell capacitance (pA/pF) and all TCD values are reported ± SD. Thus, an increase in the GABA_A_ receptor gating (e.g. by an agonist) results in a negative value for TCD. A positive value for TCD occurs when the current shuts. The noise was analyzed in some cases and was calculated as the center-value of a Gaussian fit to values of SD of 2 ms periods sampled at 100 ms intervals. Tonic currents, noise analysis and evoked events were analyzed in Clampfit 9 and Origin (OriginLab, 2017). Spontaneous mIPSCs were analyzed in Synaptosoft (6.07) and Origin as previously detailed (Hoestgaard-Jensen et al., 2014). Synaptic events included in the average waveform were detected with a 6*RMS threshold determined in the drug period to obviate false detections due to the increased noise during Thio-THIP application. The EC_50_ value for GBZ effect on tonic current was determined as the concentration of half-maximal effect (x_0_) in a fit to a logistic function of the general expression = 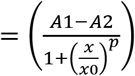.

### GABA_A_ receptor expression and FLIPR^TM^ membrane potential (FMP) Blue Assay on HEK-293 cells

The FMP assay was performed on HEK-293 Flp-In^TM^ cells expressing the human α_4_β_1_δ and α_4_β_3_δ GABA_A_ receptor as described previously (Falk-Petersen et al., 2017). In brief, HEK-293 Flp-In cells stably expressing the human GABA_A_ δ-subunit (HEK-δ cells) were transfected in a 1:1 ratio (4 μg + 4 μg) of human α_4_ (pUNIV plasmid) and human β_1_ (pUNIV plasmid) and 2:0.01 ratio (8 μg + 0.04 μg) of α_4_ to human β_3_ (pcDNA3.1) using Polyfect Transfection Reagent (Qiagen, West Sussex, UK). 16-24 hours after transfection, cells were plated into black poly-D-lysine coated clear bottom 96-well plates (BD Biosciences, Bedford, MA, USA) at 50,000 cells/well. After 16-20 hours, the media was aspirated, cells were washed and added 100 μl/well of FMP blue dye (0.5 mg/ml) (Molecular Devices (Sunnyvale, CA, USA) followed by incubation for 30 min in a humidified 5% CO_2_ incubator at 37 °C. Ligand solutions were prepared in assay buffer in 5x which for antagonist testing contained a concentration of GABA corresponding to GABA EC_80_. Ligands were added to a 96-well ligand plate and incubated for 15 min at the desired recording temperature (either 22 or 37 °C) in the NOVOstar^TM^ plate reader (BMG, LABTECH GmbH, Offenburg, Germany). The plate was read using an emission wavelength of 560 nm caused by excitation at 530 nm with response detected as changes in fluorescent signal given in fluorescent units (ΔFU). The obtained data is based on 2-3 independent experiment each with three technical replicates. All potency determinations were based on at least three independent experiments. Data was analyzed as relative changes in the fluorescent signal by taking the maximum compound induced peak/plateau signal and subtracting the baseline signal. Changes in the fluorescent signal caused by ligand addition to the wells and artefacts were manually omitted for the analysis. Concentration-response curves were fitted using the four-parameter concentration-response curve:

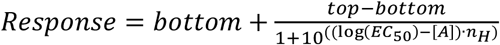

with bottom being the lower plateau response and top the upper. [A] is the logarithmic concentration of the compound and n_H_ the hill slope. Antagonists were fitted using the same equation giving IC_50_ instead of EC_50_ as for the agonist.

### Western blots

After decapitation, the cortex and hippocampus were rapidly dissected, snap-frozen on dry ice and stored at − 80 °C until use. Crude synaptically enriched membranes were prepared by quickly thawing the brain tissue at 37 °C and adding 5x w/v of 50 mM Tris HCl, pH 7.4 supplemented with EDTA-free protease inhibitors (Roche). Tissue was homogenized using 2 x 1 mm zirkonium-oxide beads in the Bullet Blender (Next Advance, NY, USA) for 30 sec at max. speed. Protein concentration was determined using Bradford protein assay and 10 μg of each sample was loaded. Blots were probed with anti-α_1_, anti-α_4_, anti-β_2_ (all a gift from Assoc. Prof. Dr. Petra Scholze, Medical University of Vienna, Center for Brain Research), anti-δ (#868A-GDN, Phosphosolutions, USA), anti-γ_2_ (#NB300-190, Novus Biologicals, Littleton, USA), anti-β3 (#NBP1-47613, Novus Biologicals), anti-GAPDH (# NB600-502, Novus Biologicals, Littleton, USA) and anti-Na^+^/K^+^-ATPase (#ab76020. Abcam, Cambridge, USA). The antibodies used are associated with the following IDs in the research resource identifier database (RRID, https://scicrunch.org/resources): AB_2631037 (GABA_A_ delta), AB_2294344 (GABA_A_ gamma2), AB_10010570 (GABA_A_ beta 3), AB_10077682 (GAPDH) and AB_1310695 (Na^+^/K^+^-ATPase). The GAPDH or Na^+^/K^+^-ATPase signal was used to normalize for loading differences. Primary antibodies were detected with HRP-conjugated goat anti-rabbit or -mouse antibody (#PI-1000 and #PI-2009, Vector Laboratories, USA) and visualized by chemiluminescence (Amersham ECL Prime). Protein levels were compared by densitometric measurement of band intensities and analyzed using Student´s t-test in GraphPad Prism 7 (GraphPad Software, San Diego, CA, USA).

### Experimental design and statistical analysis

Hypothesis testing was performed as described in the “Results” section using either one-way ANOVA with appropriate post-hoc test for multiple comparisons or t-tests for pair-wise comparisons, depending upon the experimental design as specified in the text. In connection with each experiment the statistical test employed along with the F value (when applicable), P value, standard error and number of experiments are stated. Data are generally stated as mean ±SD except for mean potencies which are stated as pEC_50_/pIC_50_ ±SEM. For overview, number of experiments and significance levels are further provided in the figure legends.

## Results

We set out to determine the magnitude of the constitutive and the agonist-induced tonic current in DGGCs under varying conditions of temperature, free intracellular Ca^2+^ concentration and PKC activity. We found that the magnitude of the endogenous tonic current after blocking GABA_A_ receptors with 20 μM BIC differed significantly among conditions for the experiment (condition ***a*** through ***d***, figure 1A, table 1).

**Figure 1.**
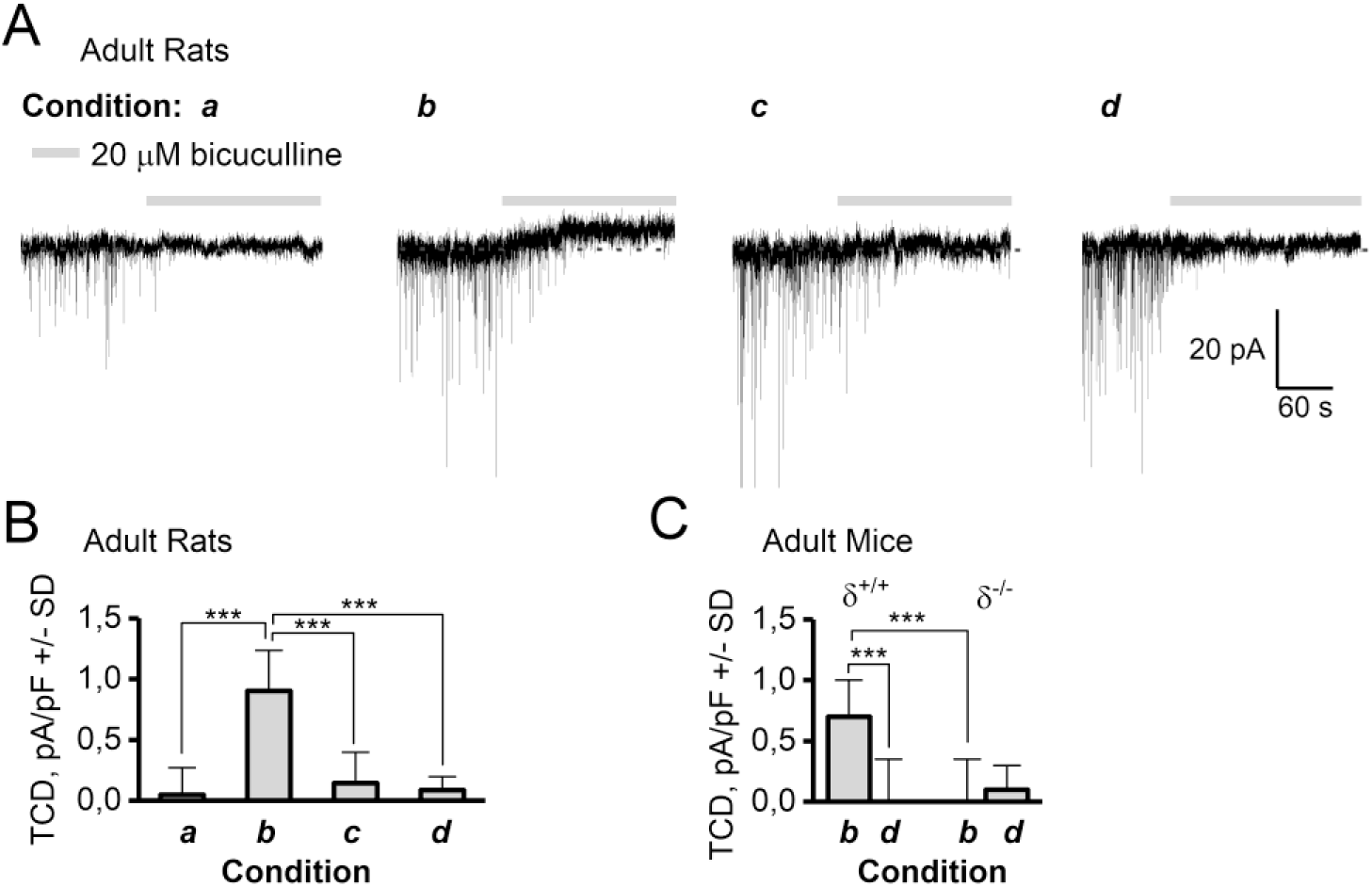
The magnitude of the GABA_A_ receptor TCD in adult rat and mouse DGGCs depend on recording condition and expression of the δ-subunit. The four recording conditions were ***a***: 24 °C and 50 μM EGTA in the ICS, ***b***: 34 °C and 50 μM EGTA in the ICS, ***c***: 34 °C including 10 μM PKC 19-36 autoinhibitory peptide and 50 μM EGTA in the ICS, and ***d***: 34 °C and 10 mM EGTA in the ICS. **A**: Representative recording traces from adult rats showing that 20 μM BIC blocks IPSCs under all recording conditions, but also blocks a tonic current in condition ***b***, which is absent in condition ***a***, ***c*** and ***d***. **B**: Summary of data from A, (n=12-14, ***: P < 0.001, ANOVA/Bonferroni). **C**: Summary of TCD in DGGCs recorded in condition ***b*** or ***d*** in δ^+/+^ and δ^-/-^ mice (n=10/11). ***: P < 0.001, paired t-test (same cell) or t-test (different cell).

**Table 1.** Summary of the effect of BIC, Thio-THIP, THIP and GBZ on the TCD in adult male rat DGGCs recorded under four different conditions (***a*-*d***). Reported values are the tonic current density (TCD), calculated as the ratio of the difference in holding current between baseline and drug period over cell capacitance. Statistical difference is determined between compound effects in the different conditions by ANOVA followed by Bonferroni mean comparison. **: P<0.01 ***: P<0.001.

|  | **Condition *a*** | **Condition *b*** | **Condition *c*** | **Condition *d*** |
| --- | --- | --- | --- | --- |
| Temperature and ICS | 24 °C | 34 °C | 34 °C, 50 μM EGTA, | 34 °C |
|  | 50 μM EGTA | 50 μM EGTA | 10 μM PKC inh. | 10 mM EGTA |
| Drug | **Tonic Current Density (TCD), pA/pF ± SD (no of cells)** |  |  |  |
| 20 μM BIC (fig.1) | 0.0 ± 0.2 (12) | 0.7 ± 0.2 *** (13) | 0.1±0.4 (14) | 0.1±0.3 (12) |
| 100 μM Thio-THIP (fig. 2) | -1.2 ± 0.4 (12) | -0.3 ± 0.3 *** (14) | -1.1 ± 0.3 (14) | -1.2 ± 0.2 (12) |
| + 100 μM PTX | -0.1 ± 0.4 | 0.9 ± 0.3 *** | 0.1 ± 0.4 | -0.0 ± 0.2 |
| 2 μM THIP (fig. 2) | -4.4 ± 0.7 (8) | -2.4 ± 0.7 ** (8) | -5.5 ± 1.0 (8) | -4.9 ± 1.2 (10) |
| + 100 μM PTX | 0.1 ± 0.3 | 0.8 ± 0.4 ** | -0.1 ± 0.3 | 0.1 ± 0.5 |
| 10 μM GBZ (fig. 3) | -1.0 ± 0.3 (10) | -0.2 ± 0.2 *** (12) | -1.1 ± 0.3 (12) | -1.2 ± 0.3 (12) |
| + 100 μM PTX | 0.0 ± 0.4 | 0.9 ± 0.3 *** | 0.2 ± 0.3 | 0.2 ± 0.2 |

Specifically, the GABA_A_ receptor-mediated tonic current density (TCD) measured using 20 μM BIC in condition ***a***, ***c*** and ***d*** was (mean pA/pF±SD) 0.0±0.2, 0.1±0.4 and 0.1±0.3, respectively, but 0.8±0.2 in condition ***b*** (one-way ANOVA F=19.5 P=1.5E-8, Bonferroni for ***b*** vs ***a***, ***c*** and ***d*** gave P<3E-6, P=1 for other combinations, n = 12-14 /condition, see table 1). We also assessed the BIC sensitive TCD in δ^-/-^ mice by recordings made in condition ***b*** and ***d*** (figure 1C). In recording condition ***b***, a TCD mean pA/pF±SD of 0.7±0.3 was measured in δ^+/+^ mice compared to 0.0±0.4 in δ^-/-^ mice (P<0.01, unpaired t-test, n= 9/10). In recording condition ***d***, the BIC-induced TCD was 0.0±0.4 in δ^+/+^ and 0.1±0.2 in δ^-/-^ mice (ns, n=10/11), indicating that the receptors responsible for the Ca^2+^, PKC and BIC sensitive TCD are dependent on incorporation of the δ-subunit. Because network levels of extracellular GABA were unlikely to be influenced by the composition of the ICS in a single neuron, it appears plausible that the TCD in condition ***b*** (figure 1A, B) is a result of predominantly constitutively active receptors, thus inferring BIC as an inverse agonist.

To proceed, two cases were considered: If extracellular levels of GABA mediated the TCD in condition ***b***, GABA_A_ receptors could be internalized in conditions ***a***, ***c*** and ***d***, and thus not in the capacity to respond to the extracellular levels of GABA. If, on the other hand, the TCD in condition ***b*** was due to constitutive activity, then channels must be either shut or internalized in conditions ***a***, ***c*** and ***d***. In either case, the effect of bath-applied orthosteric agonists should reveal if the recording conditions had created a functional difference in the number of receptors available for activation. We therefore tested the δ-subunit selective agonists Thio-THIP (100 μM) and THIP (2 μM) in condition ***a*** through ***d*** (figure 2). In order to compare with effects of GBZ (below), for these and the following experiments, we used PTX rather than BIC to block agonist-induced currents.

**Figure 2.**
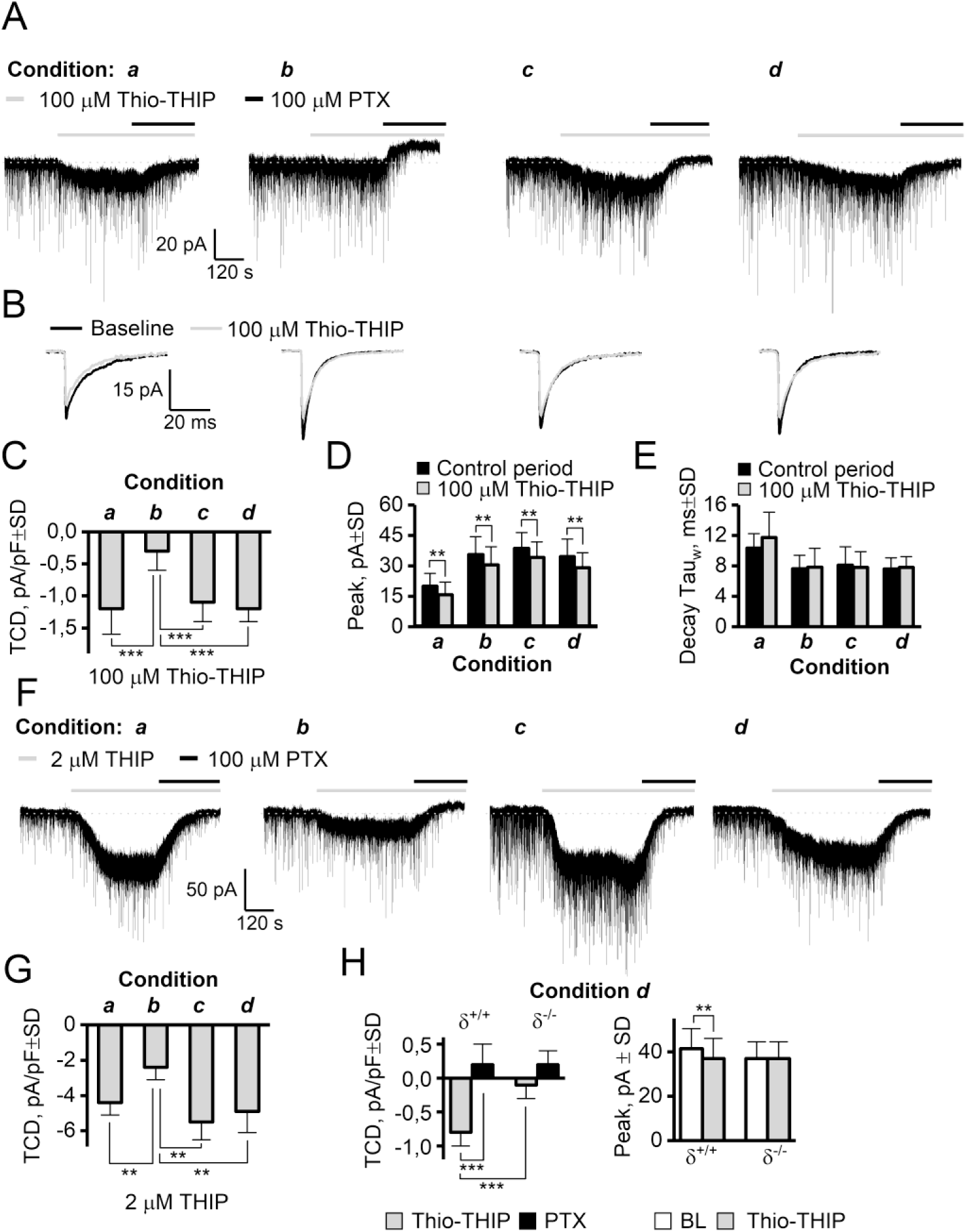
The effect of two GABA agonists, THIP (δ-subunit preferring) and Thio-THIP (α_4_β_1/3_δ selective) on the TCD is dependent on temperature, intracellular Ca^2+^-EGTA chelation and PKC activity. **A**: Full recording traces of voltage clamped (−70 mV) DGGCs upon bath application of 100 μM Thio-THIP followed by PTX. Thio-THIP induces a ~4 x larger TCD in condition ***a***, ***c*** and ***d*** than condition ***b***. Note the presence of a positive endogenous TCD in condition ***b*** only. **B**: Traces of mIPSCs display the Thio-THIP induced reduction of the average mIPSC in all recording conditions. **C**: Summary barplot of TCD values induced by Thio-THIP (n=12-14, *** P<0.001 ANOVA/Bonferroni). **D-E**: Summary of peak and decay time of the average mIPSCs (** P<0.01 paired t-test). **F-G**: Representative traces and barplot showing the effect of THIP in recording condition ***a*** through ***d*** (n=8-10, ** P<0.01 ANOVA/Bonferroni). **H**: The TCD induced by Thio-THIP in adult δ^+/+^ and δ^-/-^ mice and the effect on the average mIPSC peak in the same recordings (n =, *** P<0.001, ** P<0.01 paired t-test for same-cell measurements, unpaired for different cells).

These experiments revealed that the recording condition greatly influenced the magnitude of TCD induced by δ-subunit selective agonists in DGGCs (figure 2A, C, F-H and table 1). For 100 μM Thio-THIP recorded under conditions ***a***, ***c*** and ***d***, (figure 2A, C) the TCD was (mean pA/pF±SD) −1.2±0.4, −1.1±0.4 and −1.2±0.1, respectively, but only −0.3±0.3 in condition ***b*** (One-way ANOVA F=24.1, P=6E-10, Bonferroni for ***b*** vs ***a***, ***c*** or ***d*** gave P≤2E-7, P>0.9 for all other combinations, n=12-14/condition). Similarly to the effect of BIC, PTX administered after Thio-THIP resulted in a TCD near 0 for condition ***a***, ***c***, and ***d*** but 0.9±0.3 for condition ***b*** (one-way ANOVA F=20.1 P=1E-8, Bonferroni for ***b*** vs ***a***, ***c*** and ***d*** gave P<3E-5, P>0.79 for other combinations, n = 12-14/condition, see table 1). The change in noise, a measure reflecting open-channel probability (only condition ***b*** and ***d*** for Thio-THIP analyzed), was ns from baseline to Thio-THIP in recording condition ***b*** (mean pA±SD) 1.61±0.36 and 1.65±0.39, respectively (P=0.2, paired t-test, n=14), but increased in condition ***d*** from 1.72±0.34 during baseline to 2.2±0.4 in Thio-THIP (P=0.02, paired t-test, n=12). We did not detect a significant difference in baseline noise in recording condition ***b*** vs ***d*** (P=0.4, t-test). The TCD observed for THIP (2 μM, figure 2F, G) displayed a similar effect as Thio-THIP. In this case, the effect of THIP in conditions ***a***, ***c*** and ***d*** resulted in a TCD of (mean pA/pF±SD) −4.4±0.7, −5.5±0.9 and − 4.9±1.2, respectively, but −2.4±0.7 in condition ***b*** (one-way ANOVA F=13.1, P=1E-5, Bonferroni for ***b*** vs ***a***, ***c*** and ***d*** gave P ≤ 0.007, P>0.32 for all other combinations, n=8-10/condition, see table 1). The effect of THIP was reversed to baseline levels by PTX for condition ***a***, ***c*** and ***d*** and to levels above the baseline for condition ***b*** (one-way ANOVA F=11.8 P=1E-6, Bonferroni for ***b*** vs ***a***, ***c*** and ***d*** gave P<4E-3, P>0.8 for other combinations, n = 8-10/condition, see table 1) figure 2 and table 1). Because of the significantly increased response to orthosteric agonists in conditions ***a***, ***c*** and ***d*** vs ***b***, we conclude that a net receptor internalization could not be the reason for the absent constitutive current under condition ***a***, ***c*** and ***d***. While the TCD induced by δ-subunit specific orthosteric agonists was highly dependent on recording condition (figure 2A, C-G), the peak of the averaged non-contaminated mIPSC was, however, similarly diminished by Thio-THIP in all conditions (figure 2B, 2D, E). The reduction in the synaptic mIPSC occurs because peri-synaptic α_4_β_1/3_δ receptors activated by bath applied Thio-THIP are not available for “further” activation by synaptically released GABA, and therefore the peak is composed of fewer GABA-activated channels, and thus reduced. This suggested that peri-synaptic and Thio-THIP-responding receptors contributing to the mIPSC are not subject to the same regulation imposed by the recording conditions as extrasynaptic receptors, and probably not similarly phosphorylated as non-synaptic receptors. For Thio-THIP (figure 2D), the peak mIPSC was reduced from (mean pA±SD) −20.0±6.3 to − 15.8±6.2 (condition ***a***, n=12, P=2E-6, paired t-test), in condition ***b*** from −35.6±8.8 to −30.4±9.0 (n=14, P=2E-6), in condition ***c*** from −38.7±7.6 to −34.1±7.6 (n=14, P=0.01) and in condition ***d*** from −34.6±8.7 to −29.1±7.4 (n=12, P=0.001). The selectivity of Thio-THIP for δ-subunit containing receptors in mediating effects on TCD and mIPSC peak was assessed in δ^-/-^ mice (figure 2H). Recording from DGGCs in condition ***d***, Thio-THIP (100 μM) induced a TCD of (mean pA/pF±SD) −0.8±0.2 in δ^+/+^ mice, but −0.1±0.2 in δ^-/-^ mice (t-test, n=9/10, P =8E-7). Similarly, the Thio-THIP-induced reduction in mIPSC peak, observed in rat DGGCs (figure 2B, D) was also seen in δ^+/+^ mice (−41.4±9.0 pA in baseline vs −37.4±9.0 pA in Thio-THIP, paired t-test, n=9, P=0.01), but absent in δ^-/-^ mice (−37.1±7.6 in baseline vs −37.2±7.5 in Thio-THIP, paired t-test, n=10, P=0.9). We did not analyze the synaptic response to THIP, because the noise associated with the effect led to very significant distortion of the waveform, interfering with detection and analysis.

It is commonly presumed that GBZ cannot close GABA_A_ receptor channels displaying constitutive activity (Bai et al., 2001; McCartney et al., 2007; Wlodarczyk et al., 2013). Therefore, the recognized ability to minimize the constitutive gating through the recording condition also offered a new and untried possibility for probing the antagonist effects of GBZ. If GBZ is a neutral antagonist at constitutively active receptors, it follows that we should expect a TCD near zero for GBZ in the conditions under which the constitutive gating is shut. However, in these conditions, GBZ (10 μM), although completely blocking all mIPSCs, also induced a TCD, i.e. an apparent receptor activation, of (mean pA/pF±SD) −1.0±0.3, −1.1±0.3 and −1.2±0.3 in condition ***a***, ***c*** and ***d*** respectively, but only −0.2±0.2 in condition ***b*** (one-way ANOVA, F=20.7, P=2E-8, Bonferroni for ***b*** vs ***a***, ***c*** and ***d*** gave P<1E-5, P>0.62 for other combinations, n=10-12/condition). The effect of GBZ introducing a TCD in condition ***a***, ***c*** and ***d*** was reversed to baseline levels by 100 μM PTX, and for condition ***b***, to above baseline, producing a TCD of 0.9±0.3 pA/pF±SD (one-way ANOVA F=19.2 P=7E-8, Bonferroni for ***b*** vs ***a***, ***c*** and ***d*** gave P<3E-4, P>0.71 for other combinations, figure 3, table 1). The block of the mIPSCs by GBZ was complete in < 20 secs from onset, and coincident with onset of the TCD, which continued to develop for at least 2 min (figure 3). A concentration-response relation of GBZ for this effect recorded in condition ***d*** gave an EC_50_ of 2.1 μM (five concentrations in the range 10 nM to 50 μM GBZ tested, figure 3D, one concentration/cell). When reversing the order of administration for PTX (to 1^st^) and GBZ (2^nd^) in recording condition ***d***, GBZ produced no change in TCD (0.0±0.3 pA/pF±SD, n = 4). However, if the constitutive current in condition ***b*** was first shut by 20 μM BIC before perfusion of GBZ (10 μM), we found that GBZ now induced a TCD of −1.3±0.4 pA/pF±SD which again could be shut by 100 μM PTX (figure 3B, as originally observed in (Wlodarczyk et al., 2013). The difference in TCD produced by GBZ (10 μM) in condition ***b***+20 μM BIC (−1.3±0.4 pA/pF±SD, n= 8) and condition ***c*** or ***d*** (−1.1±0.2, −1.2±0.2 pA/pF±SD, n = 14 and 12, respectively, table 1) was non-significant (one-way ANOVA, F=1.89, P=0.16).

**Figure 3.**
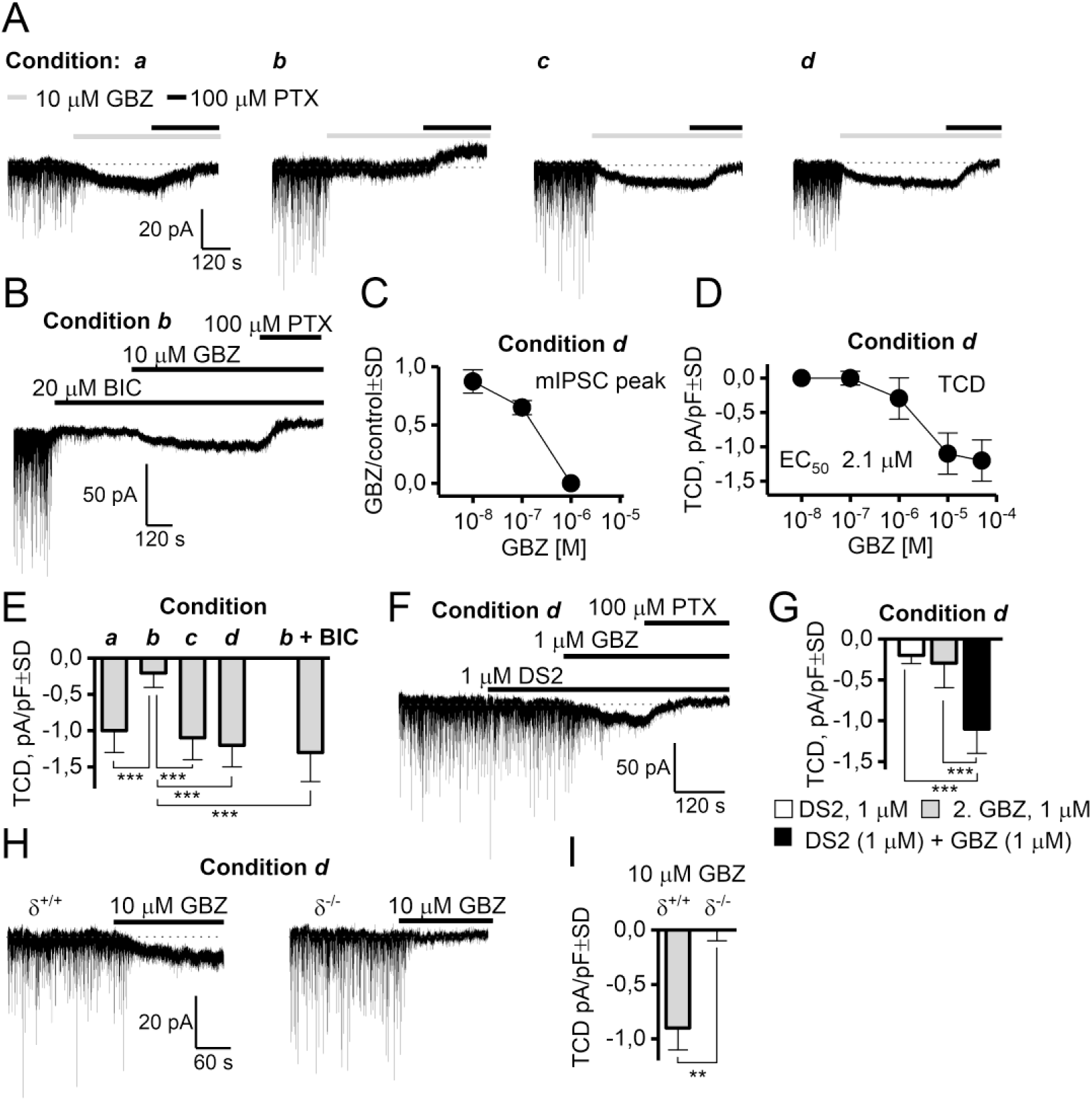
GBZ induces a δ-subunit dependent TCD in adult rat DGGCs under recording conditions phenomenologically characterized by the absence of a constitutive current. **A**: Recording traces displaying the effect of GBZ and PTX in recording conditions ***a***-***d***. GBZ quickly blocks mIPSCs, but also induces a TCD in condition ***a***, ***c*** and ***d***, which is reversed by PTX. **B**: Shutting of the constitutive current by BIC (20 μM) in recording condition ***b*** is reversed by GBZ (10 μM). **C-D**: Concentration-response relation of GBZ in blocking mIPSCs, and inducing a TCD in recording condition ***d***. **E**: Summary of effects of GBZ to induce a TCD in recording condition ***a***-***d***, incl. recording condition ***b*** in which the constitutive current is shut by BIC. **F-G**: The δ-subunit selective positive modulator DS2 potentiates the effect of GBZ in producing a PTX-sensitive tonic current. **H-I**: Recordings of DGGCs from adult δ^+/+^ and δ^-/-^ mice demonstrate that the effect of GBZ inducing a TCD is dependent on expression of the δ-subunit. ** P<0.01, t-test (figure I), *** P<0.001, ANOVA/Bonferroni.

To assess if this apparent agonistic effect of GBZ was dependent on GABA_A_ receptors incorporating the δ-subunit, we first tested GBZ (1 μM) in the presence of the δ-subunit positive modulator DS2 (1 μM). In recording condition ***d***, bath application of DS2 (1 μM) and subsequent (3 min later) administration of GBZ (1 μM) increased TCD to −1.1±0.3 pA/pF±SD (n=8). This was significantly larger than the TCD for application of 1 μM DS2 (−0.2±0.1, n=5) or 1 μM GBZ (−0.3±0.5, n=8) alone (one-way ANOVA, F=16.5, P=8.6E-5, Bonferroni P<3E-4) albeit analysis for synergism of the combination value of −1.1±0.3 vs the calculated sum (−0.5±0.3, independence assumed), was not made. Further, the effect of GBZ (10 μM) recorded under condition ***d*** in δ^-/-^ mice was 0.0±0.1, but −0.9±0.2 in δ^+/+^ mice (n =7/7, P=0.002, t-test, figure 3I), indicating that the effects of GBZ in producing a TCD was indeed dependent on presence of the δ-subunit.

Finally, we assessed whether the effect of GBZ in producing a TCD could be mediated through homomeric rho1 GABA_A_ receptors and tested 10 μM TPMPA in condition ***d***, which was ineffective in blocking the effects of GBZ (TCD was −1.1±0.3 pA/pF in GBZ vs −1.3±0.3 pA/pF in GBZ + TPMPA, P=0.4, n=5, t-test). However, despite these clear effects of GBZ in inducing a current in receptors incorporating the δ-subunit with at least partial contribution from the β_1_ subunit (below, figure 5), we could not demonstrate such effects of GBZ in HEK-293 cells expressing human recombinant α_4_β_1/3_δ receptors (figure 4). Using a recently characterized assay employing voltage-sensitive fluorescence emitting dyes (Falk-Petersen et al., 2017), the effect of GABA, GBZ and BIC was characterized at human α_4_β_1_δ receptors at 37 and 22 °C. At 37 °C, GABA displayed an EC_50_ value of 0.35 μM (pEC_50_=6.5±0.13, n=3) (figure 4A), and GBZ displayed an IC_50_ value of 0.42 μM (pIC_50_=6.4±0.20, n=3) against a concentration of EC_80_ of GABA (figure 4B). Similarly, at human α_4_β_3_δ receptors, GABA displayed an EC_50_ value of 0.35 μM (pEC_50_=6.5±0.03, n=3) (figure 4D) and GBZ displayed an IC_50_ of 0.50 μM (pIC_50_=6.3±0.07, n=3) (figure 4E). The constitutive activity at both 37 °C and 22 °C was neglible (no effect of BIC at baseline activity, not shown), and we did not observe any current induced by GBZ at neither receptor subtype (figure 4C, F).

**Figure 4.**
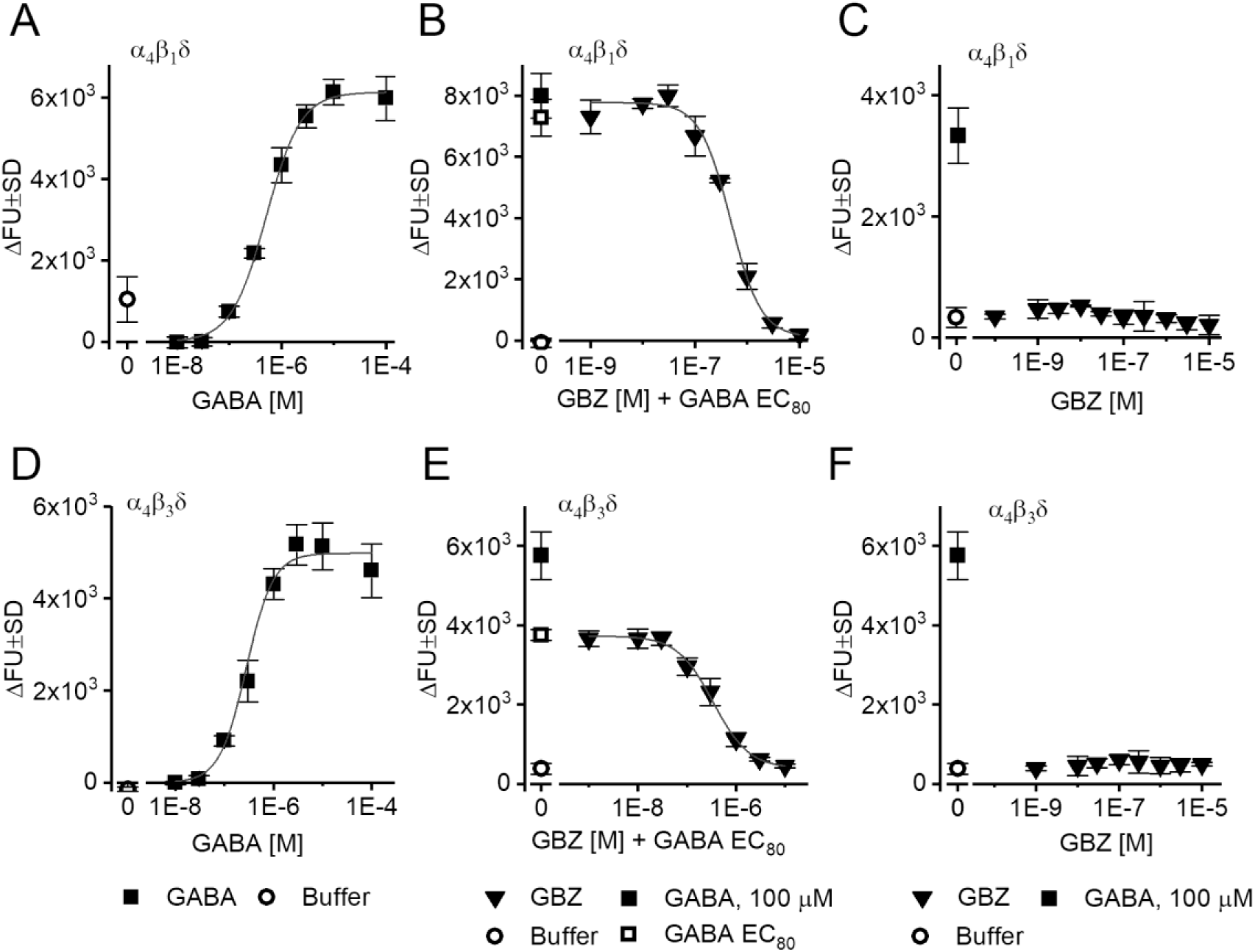
GBZ antagonizes effect of GABA but does not induce a current alone in constitutively silent α_4_β_1_δ and α_4_β_3_δ receptors expressed in HEK-293 Flp-In cells. **A, D**: GABA (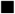) activates human α_4_β_1_δ (A) and α_4_β_3_δ (D) receptors with a similar EC_50_ of 3.5E-7 M. **B, E**: GBZ (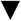) concentration-dependently blocks the response to an EC_80_ concentration of GABA with IC_50_ = 4.2E-7 M at α_4_β_1_δ receptors (B) and 5E-7 M at α_4_β_3_δ receptors (E). **C, F**: GBZ does not induce a current alone in any receptor type. All datapoints in graph **A-F** are means of triplicate measurements ±SD. FU, fluorescence unit. Symbols below belong to both figures in the three vertical panels.

**Figure 5.**
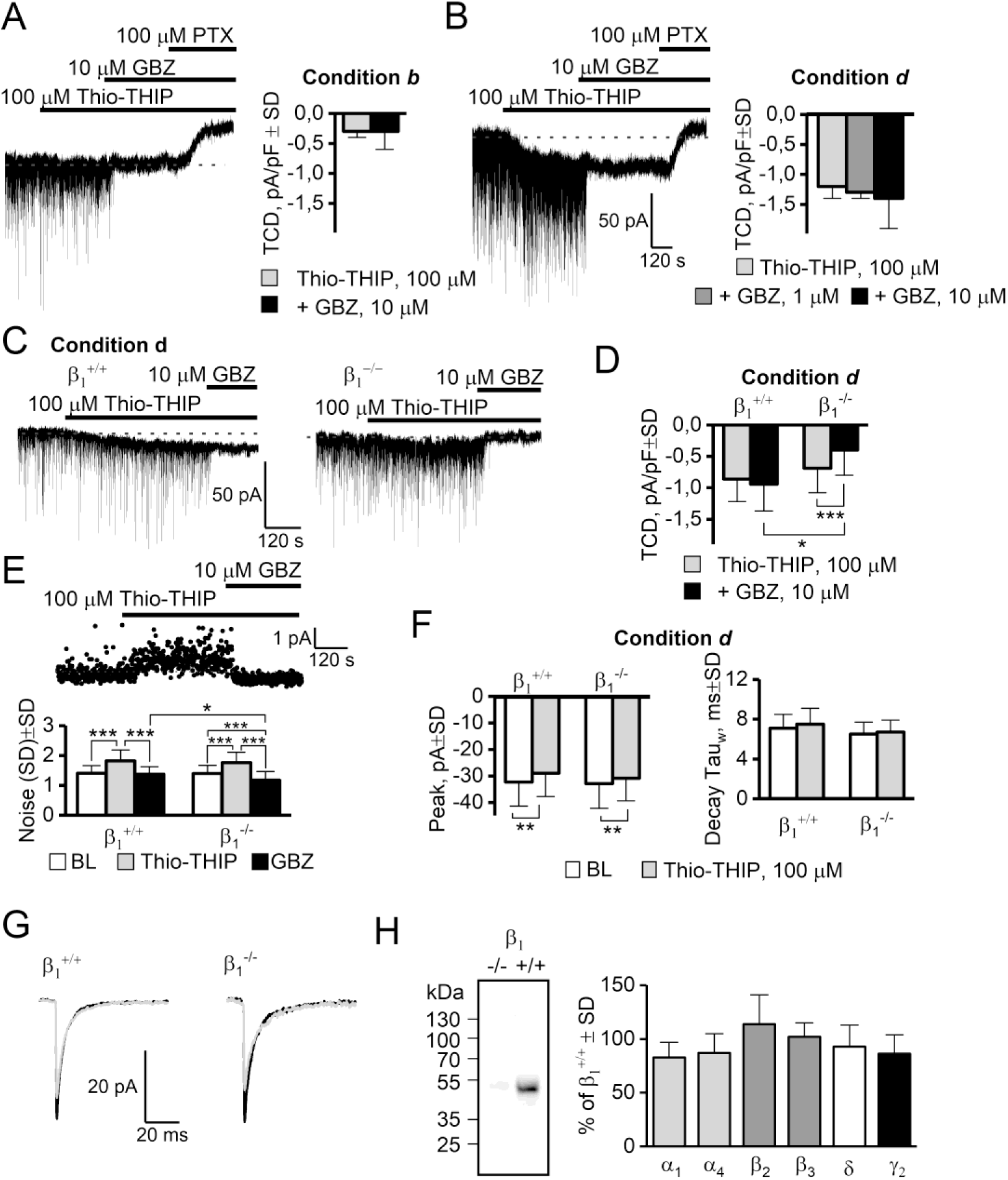
GBZ does not antagonize the effect of Thio-THIP. **A**: Thio-THIP, GBZ and PTX on DGGCs in recording condition ***b*** in adult rats (n=8). **B**: Same experiment as A, recording condition ***d***. Note that 10 μM GBZ neither blocks, nor increases the TCD already induced by Thio-THIP (right panel, n=13). **C-D**: Thio-THIP and GBZ recorded in condition ***d*** in adult β_1_^+/+^ and β_1_^-/-^ mice (n=18/22). **E**: The noise (SD) induced by Thio-THIP and GBZ in β_1_^+/+^ and β_1_^-/-^ mice. The noise in Thio-THIP + GBZ is significantly lower in β_1_^-/-^ than β_1_^+/+^ mice. **F-G**: The average mIPSC peak is similarly reduced by Thio-THIP in β_1_^+/+^ and β_1_^-/-^ mice. **H**: Western blots of whole hippocampal homogenate from β_1_^+/+^ and β_1_^-/-^ mice, displaying absence of β_1_ subunit immunoreactivity in β_1_^-/-^ mice (left) and no significant compensatory changes in α_1_, α_4_, β_2_, β_3_, δ and γ_2_ subunits in β_1_^-/-^ mice compared to β_1_^+/+^ (right, see extended data figure 5-1 for individual WBs).

Because the effect of GBZ and Thio-THIP were both dependent on the presence of the δ-subunit and both ineffective at producing a significant TCD when the constitutive gating was present (condition ***b***, figure 2 and table 1), we hypothesized that their effects were mediated by the same receptors. We tested in conditions ***b*** and ***d*** if GBZ (1 and 10 μM) could affect the TCD induced by Thio-THIP (100 μM). In condition ***b***, Thio-THIP induced a TCD of −0.3±0.3 pA/pF±SD which was not changed by 10 μM GBZ (paired t-test, P=0.6, n=8, figure 5A). In condition ***d***, the TCD induced by Thio-THIP (−1.2 ±0.2 pA/pF±SD, n = 13, figure 5B) also was not significantly changed by 1 μM GBZ (paired t-test, n=5, P=0.25) or 10 μM GBZ (paired t-test, n=8, P=0.1).

Because the effects of GBZ (at 10 μM, maximum effect, figure 5B) and Thio-THIP were not additive, this suggested that their effects could be mediated through the same (saturable) mechanism. Thio-THIP displays a selectivity for the α_4_β_1/3_δ receptor subtype, sparing other hippocampal δ-subunit partnerships (for a list of receptor configurations tested, see (Hoestgaard-Jensen et al., 2014)). To assess if a β subunit could be involved in this response, we repeated the Thio-THIP–GBZ interaction study in mice ablated of the β_1_ subunit and made recordings in condition ***d*** (figure 5C-G). In the β_1_^+/+^ mice, Thio-THIP induced a TCD of (mean pA/pF±SD) −0.9±0.3 compared to −0.7±0.4 in β_1_^-/-^ mice (n=18/22, P=0.18). However, the ensuing effect of GBZ (10 μM) produced different effects on the TCD between genotypes. In β_1_^+/+^ mice, GBZ added after Thio-THIP, produced a TCD of −0.9±0.4 (Thio-THIP vs Thio-THIP+GBZ, n=18, P=0.5 paired t-test), but significantly reduced the TCD in β_1_^-/-^ mice to −0.4±0.4 (Thio-THIP vs Thio-THIP+GBZ, n=22, P=1E-4 paired t-test). The noise (SD) during baseline and during Thio-THIP application was independent on genotype (figure 5E), whereas GBZ reduced noise in β_1_^-/-^ to a level below that of the β_1_^+/+^ (GBZ+Thio-THIP in β_1_^+/+^ produced a noise of 1.37±0.26 but 1.18±0.29 in β_1_^-/-^ mice (P=0.04, n=22/18, t-test). An additional qualitative observation was that the onset of the noise by Thio-THIP, or the shutting of it by GBZ, was fast (see figure 5A-B and 5G), reaching a plateau within tens of seconds after onset, whereas the TCD by either Thio-THIP or GBZ continued to develop for typically 2 minutes after onset. These data obtained in β_1_^-/-^ mice infer that the GBZ response involves, at least in part, β_1_-containing receptor subtypes – in addition to δ-subunit containing receptors, possibly in a common subtype, e.g. α_4_β_1_δ.

We next wanted to analyze possible compensatory changes in GABA_A_ receptor subunit levels as a consequence of the β_1_ deletion. To address this we compared the expression levels of α_1_, α_4_, β_2_, β_3_, δ and γ_2_ GABA_A_ subunits in β_1_^-/-^ and β_1_^+/+^ counterparts by Western blotting. As illustrated in figure 5H, β_1_^-/-^ and β_1_^+/+^ whole hippocampal tissue did not exhibit significant differences in the content of the subunits (blots in extended data set, figure 5-1). Albeit only the entire hippocampus was investigated, these data support that the β_1_-subunit plays a direct role in the observed effects of GBZ, and that this is not a result of compensatory mechanisms pertaining to related subunits, either by sequence or expression profile.

Despite the fact that GBZ and Thio-THIP induced a TCD of similar magnitude, GBZ did so without any increase in the noise. A functional arrangement that may explain and link these discrepant observations is by considering the possibility of a predominately dendritic location of the GBZ current source. Due to the filtering properties of the dendrites, the visible noise must originate electrotonically close to the soma (recording electrode), whereas the magnitude of the TCD, the DC component, is much less restricted by filtering (Spruston et al., 1993; Johnston, 1995). We assessed the possibility that the TCD induced by GBZ could have significant distal dendritic contribution in rats by comparing the effect of GBZ (1 μM) on evoked events from electrotonically proximal and distal loci, recorded from rat DGGCs in condition ***b*** and ***d*** (figure 6). The peak current was normalized to the slope of the I-V curve and the reversal potential for the proximal and distal events calculated from linear fits demonstrating the presence of a significant space-clamp error resulting in a difference of 21.8 mV for the reversal potential of the proximal vs the distal event (figure 6A-B). The difference in reversal potential can be used to identify the origin of the spontaneous- and mIPSCs, because spontaneous IPSCs reversed at the same potential as the proximally evoked inhibitory event (not shown), arguing for an origin of spontaneous IPSCs at electrotonically short distances from the soma (see also (Soltesz et al., 1995)). A few neurons were filled with biocytin to reveal the extent of the dendritic tree (figure 6C). The distance between the proximal and distal stimulation electrodes were on average 260 μm (range 210 – 290 μm), largest for cells recorded from slices that originated ~350 μm ventral to the hippocampal commissure (which was the most dorsal slices recorded from). The inhibitory effect of GBZ (1 μM) on proximal and distal evoked inhibitory events (figure 6D), recorded in condition ***b*** and ***d*** showed that a distal component was reduced significantly less, and at a slower rate in condition ***d*** than ***b***.

**Figure 6.**
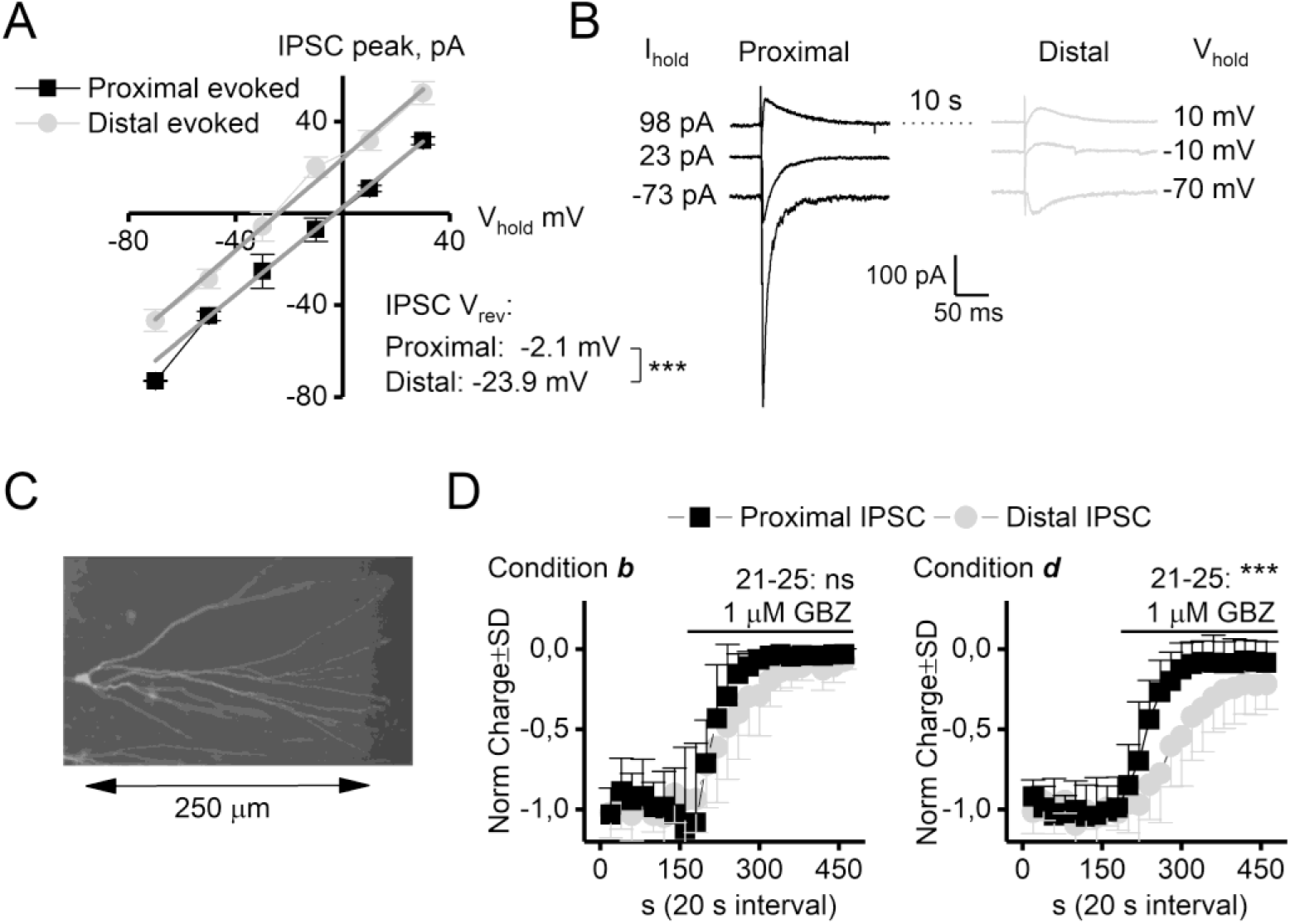
GBZ is less effective at GABA_A_ receptors at distal loci in DGGCs. **A**: IV relation of evoked events from proximal and distal loci. The results were normalized to the GABA_A_ receptor conductance for the individual neuron. The reversal potential for proximal and distal evoked events differed by 21.8 mV, indicating a considerable space clamp error. **B**: Representative traces at −70, −10 and 10 mV. **C**: Biocytin filled and stained DGGC. **D**: Normalized proximal and distal events are not equally inhibited by GBZ, yielding a slower effect of GBZ in recording condition ***d*** compared to ***b*** and demonstrating less inhibition at the average response of event 21-25. *** P< 0.001, t-test, n=9/10.

## Discussion

In the present study we show that the magnitude of the constitutive and the agonist-induced tonic current passed by possibly α_4_β_1/3_δ GABA_A_ receptors in adult rodent DGGCs is determined by temperature, free intracellular Ca^2+^ levels and PKC activity. By comparing effects of the orthosteric agonists THIP and Thio-THIP, the orthosteric antagonists BIC and GBZ, and the channel blocker PTX in four different recording conditions in adult rats, δ^+/+^, δ^-/-^, β_1_^+/+^ and β_1_^-/-^ mice, we demonstrate the existence of an inverse relationship of the constitutive vs. the agonist-induced tonic current in DGGCs. Further, we show that only under the conditions of a shut constitutive current, does GBZ induce a PTX-sensitive tonic current. This current is completely dependent on the expression of the δ- and to some degree also the β_1_-subunit, elucidated with the first *in vitro* phenotypical description of a β_1_^-/-^ mouse strain. However, this effect of GBZ could not be replicated in human α_4_β_1/3_δ receptors expressed in HEK-293 cells, suggesting a component missing in HEK cells, compared to DGGCs. Our data suggest that *in vivo* efficacy of δ-subunit specific agonists (i.e. THIP) in DGGCs, is diminished in conditions of increased PKC activity.

### Constitutive activity of GABA_A_ receptors

The four different recording conditions (***a***-***d***) were chosen because of previous findings that implicate temperature and kinase activity as regulators of δ-subunit containing GABA_A_ receptors (Wei et al., 2003; Houston et al., 2007; Bright et al., 2011; Bright and Smart, 2013). The different effect of conditions (table 1), suggest that the endogenous TCD is mediated by intracellular high activity of kinase activity in condition ***b***, most likely mediating specific phosphorylation of receptor subunits (discussed below). Recordings at 24 °C gave similar result as kinase inhibition at 34 °C, which we suggest is a consequence of low kinase activity at 24 °C, because PKC in this case does not readily translocate to the plasma membrane in mammalian cells (Chemin et al., 2007). Since the responses to the orthosteric agonists Thio-THIP and THIP were increased in condition ***a***, ***c*** and ***d*** (relative to ***b***, figure 2, table 1, see below), the same conditions for which the TCD was absent (***a***, ***c*** and ***d***, figure 1, table 1), two conclusions are made. First, the absence of an endogenous TCD in condition ***a***, ***c*** and ***d***, is not due to internalization of δ-subunit containing receptors, and second, the TCD in condition ***b*** must be the result of constitutively active receptors that can be shut by PKC inhibition, BIC and PTX (figure 1 and 2, table 1) but not by GBZ (see below). The constitutively active receptors recorded in condition ***b***, were also detected in δ^+/+^ but not in δ^-/-^ mice, identifying the δ-subunit as necessary and sufficient for the measured constitutive current.

### Response to THIP and Thio-THIP in different recording conditions

Results obtained with THIP and Thio-THIP were qualitatively similar, although the higher effect-ratio between conditions ***b*** and ***c*** or ***d*** (~ 2 for THIP but ~4 for Thio-THIP) suggest that PKC inhibition or chelating of intracellular Ca^2+^ affect receptors that are targets for Thio-THIP more selectively. We shall therefore focus this discussion on α_4_β_1/3_δ receptors due to the selectivity of Thio-THIP (Hoestgaard-Jensen et al., 2014). Since the α_4_β_1/3_δ, but not α_4_β_2_δ receptors, exhibit constitutive gating in oocytes and upon PKA-activation in HEK cells (Tang et al., 2010; Jensen et al., 2013; Hoestgaard-Jensen et al., 2014), these subtypes appear plausible candidates for the constitutive gating in DGGCs. Studies examining the co-existence of a PKA activated constitutive current and GABA activated current in α_4_β_3_δ GABA_A_ receptors in HEK cells, demonstrated that the constitutive current sets the baseline floor, under which sufficiently low concentrations of agonist is ineffective at increasing channel open-probability (Tang et al., 2010). Our data in DGGCs are in support of this, since the effect of Thio-THIP on the TCD is minimal in condition ***b*** (high constitutive current), but ~4 times more effective in condition ***a***, ***c*** and ***d*** (no constitutive current, figure 1A, 2A).

The effect of a high concentration of EGTA or inhibitor of PKC in the ICS on the endogenous TCD and response to agonists, suggests an already high level of endogenous kinase activity and receptor phosphorylation at 34 °C in low concentration of EGTA in the ICS (see also (Brandon et al., 2000; Bright et al., 2011; Bright and Smart, 2013)). It is therefore interesting to note that the effect of Thio-THIP on the synaptic response was independent on the recording condition (figure 2B), suggesting that Thio-THIP-responding receptors in the synapse are not similarly sensitive to kinase activity as extrasynaptic receptors. GABA_A_ receptors interact with, among others, gephyrin, GABARAP, calreticulin (gC1q-R) and AKAP79/150 through a ~30 AA long domain in the intracellular TM3-TM4 loop (Essrich et al., 1998; Wang et al., 1999; Schaerer et al., 2001; Brandon et al., 2003). Receptor anchoring at the synapse requires gephyrin interaction, and the binding domain for gephyrin has been mapped in the β-subunits, and shown to overlap the S383 (CaMKII site) and the S408/409/410 (PKA and PKC site) (Maric et al., 2011; Kowalczyk et al., 2013; Maric et al., 2017). Trafficking of GABA_A_ and glycine receptors is sensitive to calcineurin and PKC activity due in part to phosphorylation of gephyrin (Bannai et al., 2009; Specht et al., 2011; Tyagarajan et al., 2011; Zacchi et al., 2014). However, phosphorylation of receptor subunits are likely to also play a role in anchoring to gephyrin, as phosphorylation of the S409 residue in a GABA_A_ receptor subunit β_1_ fragment also reduces the affinity to gephyrin fragments (Maric, personal communication).

### GBZ and the constitutive current in DGGCs

The two orthosteric GABA_A_ receptor antagonists, GBZ and BIC, exert different effects against spontaneously active GABA_A_ receptors: BIC shuts spontaneously active receptors and is termed an inverse agonist whereas GBZ is ineffective towards these and termed a neutral antagonist (Birnir et al., 2000; Bai et al., 2001; McCartney et al., 2007; Wlodarczyk et al., 2013). The constitutive current in recording condition ***b*** can be closed by BIC and possibly displaced by GBZ for receptors to resume constitutive gating (see (McCartney et al., 2007; Wlodarczyk et al., 2013) and figure 3B). However, since GBZ introduces a current by itself only under conditions of a shut constitutive current, we conclude that the TCD is the result of an action by GBZ at a shut receptor, and not BIC displacement. Similar to the effect of Thio-THIP in recording condition ***b*** described above, the effect of GBZ was hidden in the already active constitutive current. This effect of GBZ in DGGCs was PTX- and DS2-sensitive, completely δ-subunit dependent, TPMPA-insensitive, partially β_1_-dependent and displayed an EC_50_ of 2.1 μM. However, we were unable to demonstrate that GBZ induces a current in recombinant α_4_β_1/3_δ receptors expressed in HEK-293 cells (figure 4 A-F). It is possible that intracellular components acting through the TM3-TM4 loop at β subunits mediating the effects of GBZ in DGGCs, are missing in the HEK-293 cells. Alternatively, it is possible that δ-subunit containing α- or β-heterodimer receptors in dendrites of DGGCs with a different response to Thio-THIP, THIP and GBZ are responsible. Since GBZ had no effect on the TCD induced by Thio-THIP in β_1_^+/+^ mice, but reduced the TCD by Thio-THIP in β_1_^-/-^ mice (figure 5C, D), we conclude that, in addition to the δ-subunits, the β_1_ subunit containing receptors are also involved in the response by GBZ, but not exclusively so. We hypothesize that GBZ induces a specific receptor state, which also characterizes the phosphorylated, constitutively active α_4_β_1/3_δ receptors.

### The GBZ mediated current originates in distal dendrites

The TCD induced by GBZ in condition ***a***, ***c*** and ***d*** carried much less noise compared to the similar TCD magnitude induced by Thio-THIP in the same conditions (for mice, figure 5D, E). This discrepancy between noise and TCD may be explained by a dendritic origin of the GBZ-current, since the low-pass filtering properties of the dendrites reduce the noise, but much less the DC-component (Spruston et al., 1993; Johnston, 1995). The different response to GBZ of proximal and distal IPSCs in condition ***b*** and ***d*** (figure 6) suggests that this is indeed the case, supported by the lack of high affinity GBZ responding channels at the soma (Wlodarczyk et al., 2013) and by the identification of a temperature-sensitive population of δ-subunit containing GABA_A_ receptors in the dendrites of DGGCs (Wei et al., 2003). This very likely reflects the same population of receptors in DGGCs seen here, given that recordings at 24 °C are comparable to recordings at 34 °C including kinase inhibitors (figure 1A, 2A, 2F and 3A) with respect to effect of orthosteric agonists, constitutive current and effect of GBZ.

In conclusion, we have described the existence of kinase-dependent behavior of receptors that disguises the efficacy of orthosteric agonists and GBZ. A pathology involving increased δ-subunit containing GABA_A_ receptor phosphorylation would result in a reduced efficacy of the agonist *in vivo*, but not in a whole cell patch clamp analysis, because disease-state specific changes in intracellular Ca^2+^-homeostasis and kinase activity are, unavoidably, eliminated.

## Acknowledgments

The authors would like to thank Dr. Javier Martín from the Transgenic Core Facility at the University of Copenhagen for excellent technical assistance with generating the β_1_ KO strain. Further, we are thankful to Dr. Delia Belelli for fruitful discussions and for the generous gift of the δ mouse strain. Dr. Anders Klein is kindly acknowledged for help with animal work. This work was supported by funding from the Lundbeck Foundation (grants R133-A12270 to P.W., R192-2015-666 to U.L. and R230-2016-2562 to C.B.F.-P).

The authors declare no competing financial interests.

---

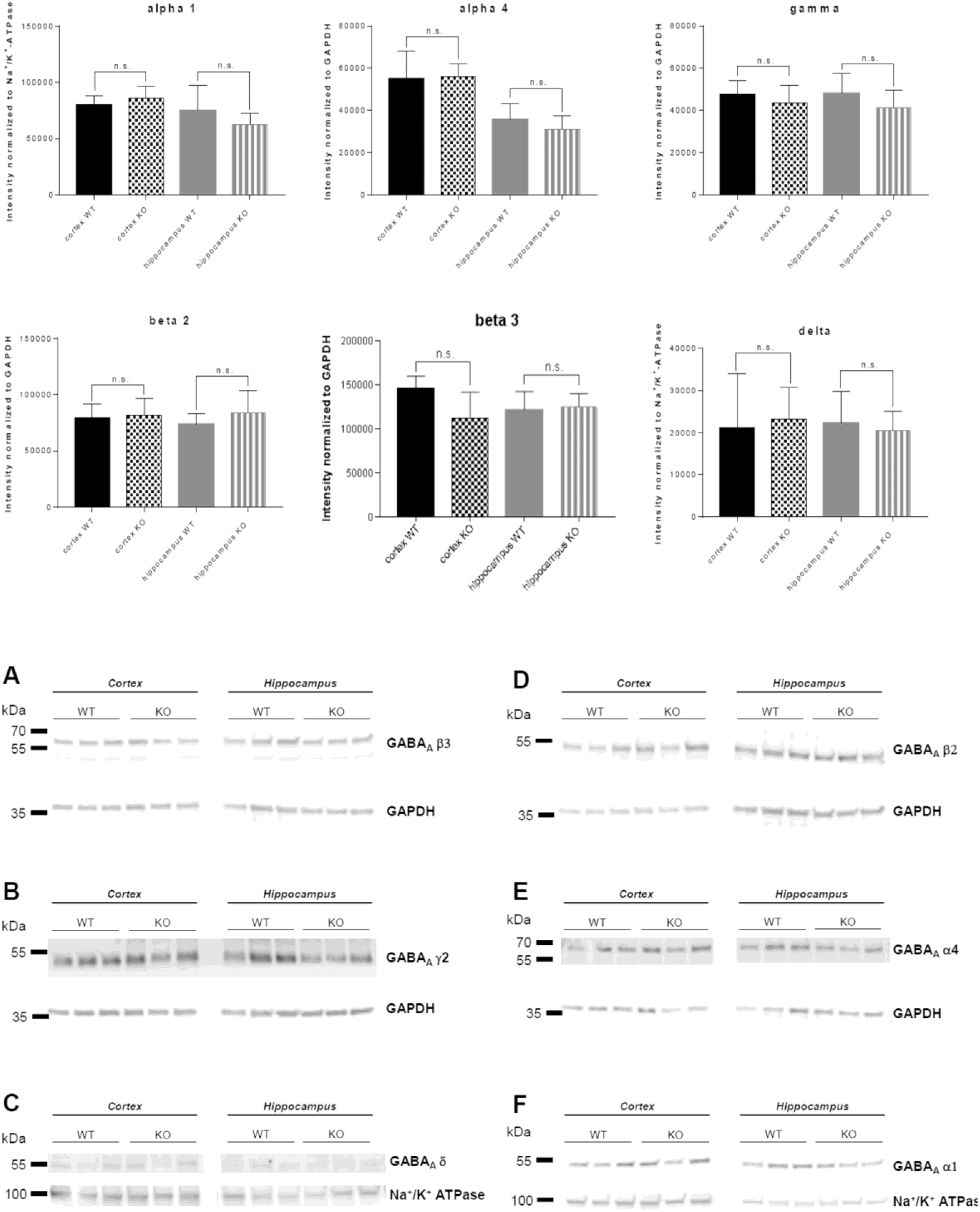

